# Low variability of dynamical functional connectivity in cerebellar networks

**DOI:** 10.1101/2020.03.16.989590

**Authors:** Izaro Fernandez-Iriondo, Antonio Jimenez-Marin, Ibai Diez, Paolo Bonifazi, Stephan P. Swinnen, Miguel A. Muñoz, Jesus M. Cortes

**Author notes:** Corresponding author (J.M. Cortes). Equal last-author contribution (Miguel A. Muñoz, Jesus M. Cortes).

## Abstract

Brain networks can be defined and explored using different types of connectivity. Here, we studied P=48 healthy participants with neuroimaging state-of-the-art techniques and analyzed the relationship between the actual structural connectivity (SC) networks (between 2514 regions of interest covering the entire brain and brainstem) and the dynamical functional connectivity (DFC) among the same regions. To do so, we focused on a combination of two metrics: the first one measures the degree of SC-DFC similarity –i.e. how much functional correlations can be explained by structural pathways– and the second one, the intrinsic variability of DFC networks across time. Overall, we found that cerebellar networks have smaller DFC variability than other networks in the cerebrum. Moreover, our results clearly evidence the internal structure of the cerebellum, which is divided in two differentiated networks, the posterior and anterior parts, the latter also being connected to the brain stem. The mechanism for keeping the DFC variability low in the posterior part of the cerebellum is consistent with another finding, namely, it exhibits the highest SC-DFC similarity among all other sub-networks, i.e. its structure constrains very strongly its dynamics. On the other hand, the anterior part of the cerebellum, which also exhibits a low level of DFC variability, has the lowest SC-DFC similarity, suggesting very different dynamical mechanisms. It is likely that its connections with the brain stem –which regulates sleep cycles, cardiac and respiratory functioning– might have a critical role in DFC variations in the anterior part. A lot is known about cerebellar networks, such as having extremely rich and complex anatomy and functionality, connecting to the brainstem, and cerebral hemispheres, and participating in a large variety of cognitive functions, such as movement control and coordination, executive function, visual-spatial cognition, language processing, and emotional regulation. However, as far as we know, our findings of low variability in the dynamical functional connectivity of cerebellar networks and its possible relation with the above functions, have not been reported so far. Further research is still needed to shed light on these findings.

## Introduction

Understanding the relationship between different classes of connectivities is pivotal in network neuroscience [1]. In particular, structural and functional connectivities are obtained from different raw data and using different pipeline analyses. Magnetic resonance imaging (MRI) provides us with structural connectivity (SC) matrices in a non-invasive manner. The entries of such matrices are given by the number of white-matter streamlines between pairs of brain regions obtained from diffusion-weighted images (DWI) combined with tractography algorithms for the reconstruction of streamlines.

MRI is also suited to construct functional connectivity (FC) matrices, that measure the similarity in the dynamics of given pairs of brain regions obtained from a sequence of functional images (accounting for the blood oxygen level dependent signals), by employing diverse metrics to characterize pairwise dynamical similarity, such as e.g. pairwise Pearson correlations or synchronization measures [2]. Here, FC networks are calculated during the resting state, a condition that is widely used for understanding the dynamics and organization of the brain basal activity in health and disease, and that is defined by the absence of any goal-directed behavior or salient stimuli [3–8].

Although SC networks and FC networks can refer to the connectivity of the same regions within the same brain, a fundamental difference between them is that both types of networks vary at very different time scales. While SC is practically invariant during the time over which FC is calculated (typically lasting for a maximum of 10 minutes), FC is well-known to vary in very short time scales, even within periods of a few seconds, exhibiting a rich dynamical repertoire (see [9] and references therein). When considering short time scales, the simplest manner to assess and quantify the temporal variations of the “dynamical functional connectivity” (DFC) is by considering a sliding window analysis. Time is divided in non-overlapping intervals of a fixed duration, and for each time window one FC is calculated; in this way the DFC is represented as a time ordered sequence of FC matrices.

Importantly, Park and Friston showed that the SC can be inferred from the DFC when the time window to calculate it is infinitely large [10]. In other words, pairwise correlations –when averaged over sufficiently long time periods– merely reflect the underlying matrix of SC (see also [11]). This fact has become even more clear with the relatively recent observation that functional networks in the resting state can be derived from the spectrum of eigenmodes –or harmonics– of the SC matrix (actually, from its associated Laplacian matrix [12]). On the other hand, for practical situations in which functional networks are obtained for much shorter time windows, how the underlying dynamics of the brain operating on a fixed SC network does generate a large repertoire of different varying functional networks, and how such dynamical patterns are related to the brain’s functionality and disease, still remain problems to be fully understood despite substantial advances achieved in the last years [10, 13–17].

The relationship between the temporally-invariant SC and the highly temporal-sensitive DFC can be assessed by comparing the two graphs at the level of individual links, a strategy which requires for symmetrical matrices *N*^2^/2 comparisons (where *N* is the number of nodes in the network). Alternatively, we follow here another and more efficient strategy that consists in establishing a comparison at a modular or community level. [18]. In particular, using a standard algorithm [19, 20], modules are identified for either structural and/or functional matrices, and this is followed by a comparison of two types of networks by using the obtained partition into modules. Our hypothesis is that, if we assume that segregated functions are associated with distinct structural modules, visualizing the functional modules in terms of the structural ones should help define and highlight how strongly structure constraints function.

In the present work, we study how SC constraints and affects DFC at the module level and find that DFC in a blindly found modules –which turns out to lie within the cerebellum– are much less variable than other networks in the cerebrum. We show two different mechanisms underlying the small variability found in cerebellar DFC, one mediated by the constraints imposed by the structural connectivity and the other one possibly related to external influences to its function. As far as we know, the small variability in cerebellar DFC networks, as observed in this study, has not been reported before and is deeply rooted in the different manner in which the cerebellum and the cerebral cortex have to deal with information processing.

## Materials and methods

### Participants

*P* = 48 healthy participants were recruited in the vicinity of Leuven and Hasselt (Belgium) from the general population by advertisements on websites, announcements at meetings and provision of flyers at visits of organizations, and public gatherings (PI: Stephan Swinnen). Participant’s age ranged between 20 and 50 years (mean age 33.9 years). None of the participants had a history of ophthalmological, neurological, psychiatric, or cardiovascular diseases potentially influencing imaging or clinical measures. All the participants provided informed consent before participation in the study, in agreement with the local ethics committee for biomedical research.

### Imaging acquisition

#### Anatomical data

A high-resolution T1 image was acquired with a 3D magnetization prepared rapid acquisition gradient echo (MPRAGE): repetition time (TR)= 2, 300 ms, echo time (TE)= 2.98 ms, voxel size = 1 × 1 × 1.1 mm^3^, slice thickness = 1.1 mm, field of view (FOV)= 256 × 240 mm^2^, 160 contiguous sagittal slices covering the entire brain and brainstem.

#### Diffusion weighted imaging

A DWI SE-EPI (diffusion weighted single shot spin-echo echo-planar imaging [EPI]) sequence was acquired with the following parameters: TR = 8, 000 ms, TE= 91 ms, voxel size = 2.2 × 2.2 × 2.2 mm^3^, slice thickness = 2.2 mm, FOV = 212 × 212 mm^2^, 60 contiguous sagittal slices covering the entire brain and brainstem. A diffusion gradient was applied along 64 non-collinear directions with a *b* value of 1, 000 s/mm^2^. Additionally, one set of images was acquired without diffusion weighting (*b* = 0 s/mm^2^).

#### Resting state functional data

Acquired with a gradient EPI sequence over a 10 min session using the following parameters: 200 whole-brain volumes with TR/TE = 3, 000/30 ms, flip angle = 90°, inter-slice gap = 0.28 mm, voxel size = 2.5 × 3 × 2.5 mm^3^, 80 × 80 matrix, slice thickness = 2.8 mm, 50 oblique axial slices, interleaved in descending order.

### Imaging preprocessing

#### Diffusion images

We applied a pre-processing pipeline similar to previous work [21–28] using FSL (FMRIB Software Library v5.0) and the Diffusion Toolkit. First, an eddy current correction was applied to overcome the artifacts produced by variation in the direction of the gradient fields of the MR scanner, together with the artifacts produced by head motion. In particular, the participant’s head motion was extracted from the transformation applied at the step of eddy current correction. The motion information was also used to correct the gradient directions prior to the tensor estimation. Next, using the corrected data, a local fitting of the diffusion tensor per voxel was obtained using the *dtifit* tool incorporated in FSL. Next, a fiber assignment by a continuous tracking algorithm was applied [29]. We then computed the transformation from the Montreal Neurological Institute (MNI) space to the individual-participant diffusion space and chose the network nodes for the calculation of SC by using a functional partition (see below).

#### Functional images

We applied a pre-processing pipeline similar to previous work [21–24, 30, 31] using FSL and AFNI (http://afni.nimh.nih.gov/afni/). First, the slice-time correction was applied to the fMRI data set. Next, each volume was aligned to the middle volume to correct for head motion artifacts. After intensity normalization, we regressed out the motion time courses, the average cerebrospinal fluid (CSF) signal and the average white matter signal. Next, a bandpass filter was applied between 0.01 and 0.08 Hz [32]. Next, the preprocessed functional data were spatially normalized to the MNI152 brain template, with a voxel size of 3 × 3 × 3 mm^3^. Next, all voxels were spatially smoothed with a 6 mm full width at half maximum isotropic Gaussian kernel. Finally, in addition to head motion correction, we performed scrubbing, by which time points with frame-wise displacement *>* 0.5 were interpolated by a cubic spline [33]. We finally removed the effect of head motion using the global frame displacement as a covariate.

### Network nodes

Both SC and DFC were built using *N* = 2514 regions of interest, thus generating networks of size *N*^2^. These 2514 regions were identified after running a method for unsupervised clustering to the functional data at the voxel level [34]. On average, each cluster –that in our study coincides with one node in the network– contained about 20 voxels of size 3×3×3 mm^3^. The algorithm takes as the input the number of desired (2514 in this case) clusters, and finds the optimal clustering solution by maximizing both within-cluster similarity and between-cluster differences while spatially constraining contiguous voxels to belong to the same cluster [34].

### Calculation of SC and DFC matrices

One SC matrix of dimension 2514 × 2514 was obtained for each participant by counting the number of white matter streamlines connecting all possible 2514 × 2514 pairs of nodes. Thus, the element matrix (*i, j*) of SC is given by the streamlines number between nodes *i* and *j*, with *i, j* = 1, …, *N* and, given the lack of directionality of streamlines, the SC is a symmetric matrix. To calculate an averaged SC matrix at the population level, we first binarized individual SC matrices and then took the overall average over participants.

With respect to functional networks, after averaging over all voxel time series within each network node, we extracted a single time series for each of the 2514 nodes. Then, DFC_*w*_ matrices were calculated by assessing the pairwise Pearson correlation coefficient between all-node time series within a time window *w*. When dividing the total time series length *T* in *W* non-overlapping windows of length *δ*, we obtained *P* × *W* matrices, where *P* is the number of participants and *W* the number of time windows. Population DFC_*w*_ matrices were built by averaging over all participants.

Remark the following considerations: DFC is a tensor, composed of a temporal sequence of squared matrices DFC_*w*_, each one with dimension *N* × *N* and calculated over a fixed window *w*. But we also can refer the two objects DFC and DFC_*w*_ at the module level, simply extracting from them the within module contributions, that we will denote respectively as DFC^*m*^ and 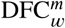, the former being another tensor composed of a sequence of matrices of dimensions *N*_*m*_ × *N*_*m*_, and the latter being a squared matrix with dimention *N*_*m*_ × *N*_*m*_ For both cases, it holds that 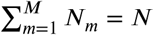.

### Adapting high-pass filtering for very short time windows

Conventional FC approaches typically work with long-time series. However, for relatively short time windows, the calculation of the matrix DFC_*w*_ requires an adaptation of the lowest bound of (LB) used for bandpass filtering of the time series by using the equation LB^−1^ = *δ* × TR, where *δ* is the window length [35]. For example, for *δ* = 10 and TR=3 seconds (used here), the high-pass filter has to be adapted by taking LB ≈ 0.03, rather than 0.01 (which was the one used here in the pipeline for the pre-processing of functional data).

### Structural modules used as a template for reordering DFC

By maximizing the modularity *Q* of the population SC matrix by employing the algorithm described in [36], we obtained a subdivision of the structural network into *M* non-overlapping modules in a way that maximizes the number of within-module links and minimizes the number of between-module links [20]. Next, we used such a partition to reorder the elements of the functional matrices and assess the link-to-link –or pairwise– similarity between SC and DFC_*w*_ matrices. This comparison was repeated for all functional matrices obtained at different windows. In this way, we have a common structural organization of modules, and this is used as a template for reordering all functional matrices^1^. As we found before [18], this is a very convenient strategy to highlight similarities and differences between both types of networks at a “mesoscopic” or community level.

### Assessment of SC-DFC similarity module by module

After reordering all functional networks using the structural modules, the SC-DFC similarity was assessed module by module, by looking at within-module links. Therefore, from the original SC and DFC_*w*_ matrices with dimensions *N* ×*N*, we extracted *m* = 1, …, *M* squared matrices SC^*m*^ and 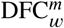 with dimensions *N*_*m*_ × *N*_*m*_, such that 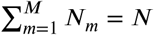. Next, for each module and window *w*, we calculate as a similarity measure the Pearson correlation 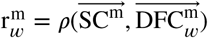, where 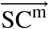 and 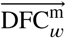 represent respectively the vector-wise representation of matrices SC^*m*^ and 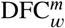. We finally averaged over windows of same size to represent the average similarity, ie., 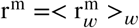.

### Assessment of DFC variations module by module

For each given window length, we obtained a series of consecutive DFC_*w*_ matrices and reordered all of them using the *M* structural modules. For each of the *m* = 1, …, *M* modules and window *w* = 1, …, *W*, we calculated the matrices 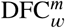, representing the within-module *m* of matrix DFC_*w*_, and assessed their variability along different time windows by calculating their pairwise spectral distance [37]:

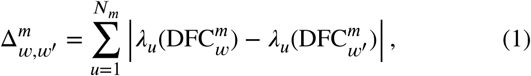

where *w, w*′ = 1, …, *W* are two generic windows, 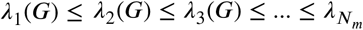 (*G*) are sets of the eigenvalues of the two graphs 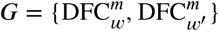, and *N* _*m*_ is the number of network nodes contained in the module *m*.

## Results

A population of young *P* = 48 healthy participants was used for the study. Diffusion and resting-state images were acquired and analyzed for each participant following the pipeline shown in figure 1.

**Figure 1:**
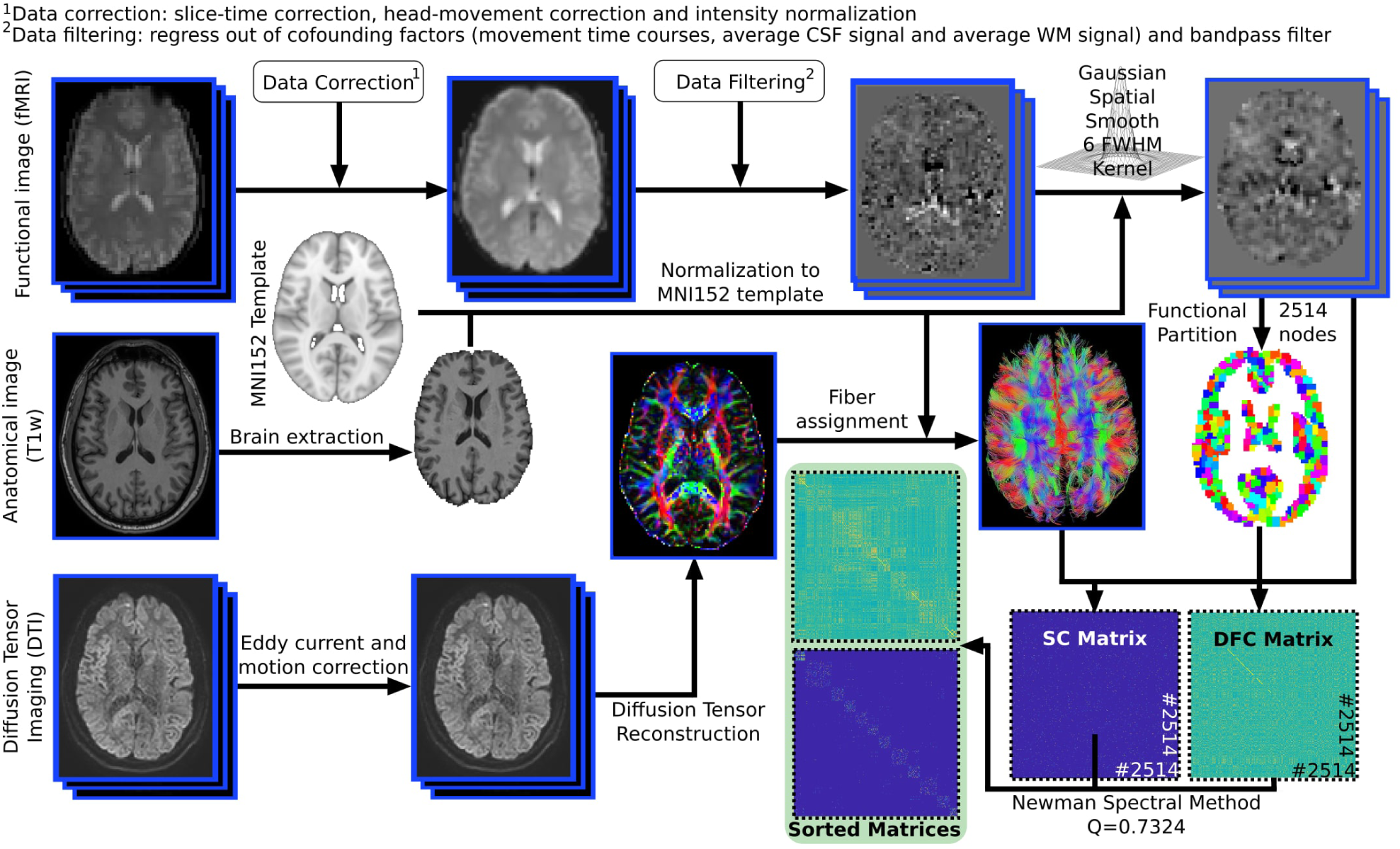
Neuroimage preprocessing pipeline. Triple acquisitions were performed per each participant: High-resolution anatomical images (T1), functional images at rest (fMRI) and diffusion tensor imaging (DTI). Following a state-of-the-art pipeline of neuroimaging preprocessing, we obtained time series of the blood oxygenation level-dependent (BOLD) signal for each network node and number of streamlines between pairs of nodes. Finally, for the comparison of SC and DFC, we re-ordered DFC according to the results after modularizing SC. Here, DFC refers to any generic window length.

First, we performed a division of the population SC matrix through modularity maximization. This resulted in M=14 non-overlapping modules (illustrated in Figure 2), with a modularity index equal to *Q* = 0.7324. Next, after calculating one DFC _*w*_ matrix per time window *w* we reordered all of them according to the optimal SC partition and calculated, for all modules separately, 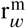, i.e. the link-to-link correlation as a measure for the similarity between SC^*m*^ and the different 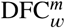, one per time window (figure 2). In particular, we first considered *δ* = 8 and averaged the measurement across all the DFC_*w*_ for the different time windows (see caption of Fig.3). Moreover, we also studied, the variability in DFC_*w*_ matrices across time windows; to this aim we measured the pairwise spectral distasnce 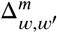 within each module *m* and time windows *w* and *w*′, and his was averaged over all pairs (*w, w*′). As illustrated in Figure 3 the analysis of SC-DFC similarity across the different modules revealed the existence of an outlier, module 1, which had a similarity value of *r*^*m*^ *>* 0.7, in contrast to the rest of modules whose similarity was lower than 0.55. The anatomy of module 1 is shown in table 1; remarkably, it is formed majorly by posterior cerebellar structures. On the other extreme, the lowest SC-DFC similarity was reported for module 14, with *r*^*m*^ *<* 0.4 for all time-window lengths. Importantly, as shown in Table 1, this module is also formed by another part of the cerebellum (its anterior part), together with other structures such as the brain stem, the fusiform and part of the lingual cortex.

**Table 1.** Brain anatomy for the three most relevant modules. Module 1, formed by posterior cerebellar structures, had the highest SC-DFC similarity independently of windows lengths. Module 11 was one of the modules with higher DFC variability along different windows, formed by several cortical regions. Module 14, with a relative DFC variability across time windows, provided the lowest SC-DFC similarity independently of window lengths and included anterior cerebellar structures, brain stem and cortical regions. For the three modules, we only reported overlapping indices bigger than 5%.

| Module 1 | Module 11 | Module 14 |
| --- | --- | --- |
| Cerebelum_Crus1 (%21.879) | Calcarine_L (%19.2356) | BrainStem (% 15.718) |
| Cerebelum_8 (%19.1888) | Temporal_Mid_R (%15.601) | Cerebelum_6 (%15.2804) |
| Cerebelum_Crus2 (%15.4499) | Lingual (%14.3722) | Fusiform (%14.2614) |
| Cerebelum_9 (%7.8111) | Temporal_Inf_R (%10.07) | Cerebelum_4_5 (%14.031) |
| Cerebelum_6 (%7.0914) | Precuneus (%8.6727) | Lingual (%10.1075) |
|  | Cuneus (%6.1875) |  |

**Figure 2:**
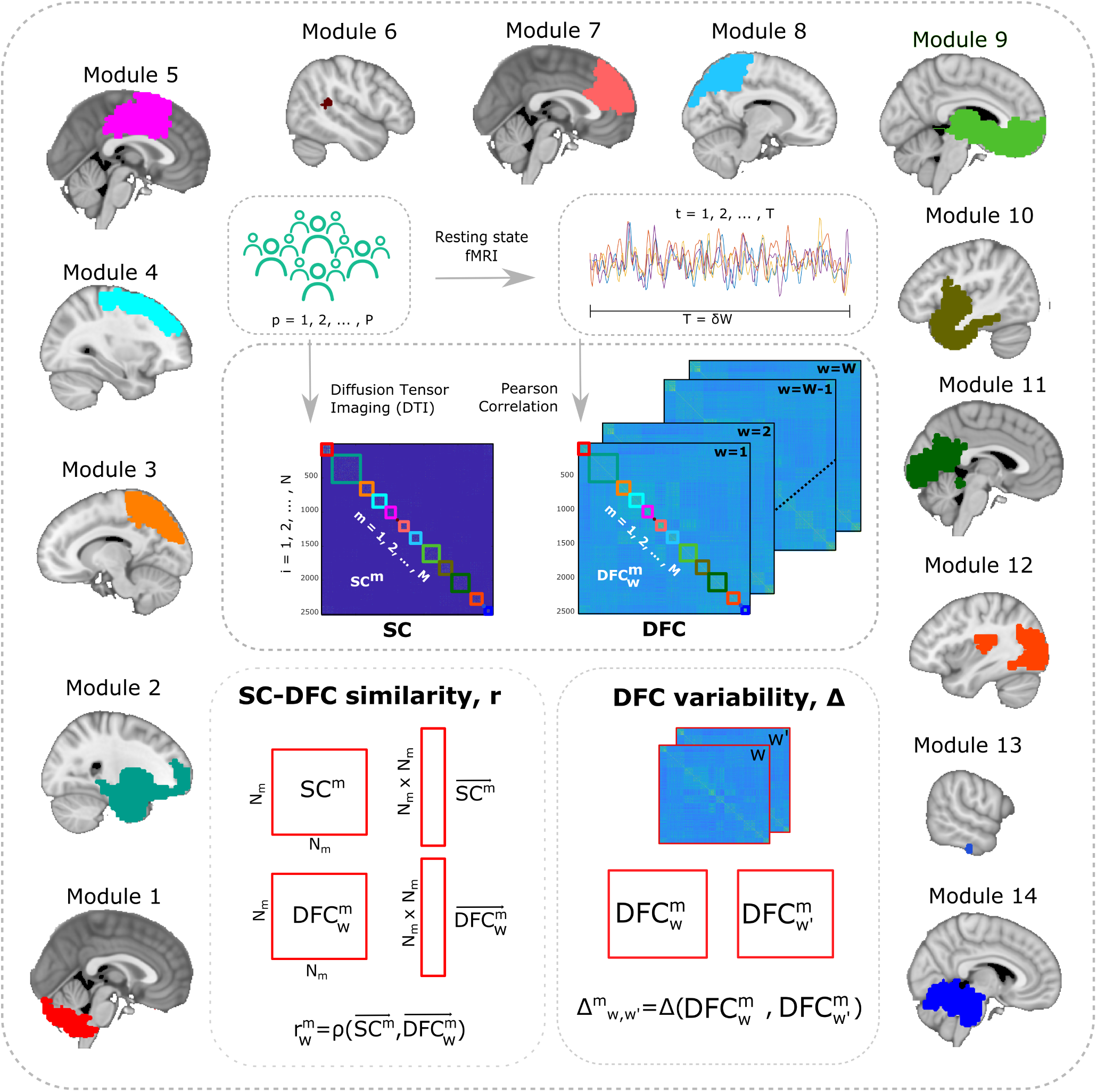
Sketch for assessing SC-DFC similarity and DFC variability module by module. Per each participant *p*, we obtained one SC matrix and a sequence of *W* matrices of DFC_*w*_. After maximization of modularity in the SC, we obtained M=14 modules, marked with squares at different colors and represented in the brain with the most representative slice for each one. All DFC_*w*_ matrices were reordered using the structural modules. Next, the SC-DFC similarity was approached within each module *m* separately, by calculating the Pearson correlation between vectorwise representations of matrices SC^*m*^ and 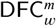. The DFC variability along different time windows was assessed by calculating the pairwise spectral distance between matrices 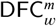 and 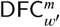. Modules 1, 5, 7, 9, 11 and 14 were bilateral; Modules 2, 4, 6, 8 and 13 were mainly located in the right hemisphere; Modules 3, 10 and 12 were mainly located in the left hemisphere. The sizes of the modules *N*_*m*_, measured in number of network nodes per module, were respectively, 176, 404, 197, 168, 180, 2, 171, 191, 251, 191, 262, 189, 1 and 131.

**Figure 3:**
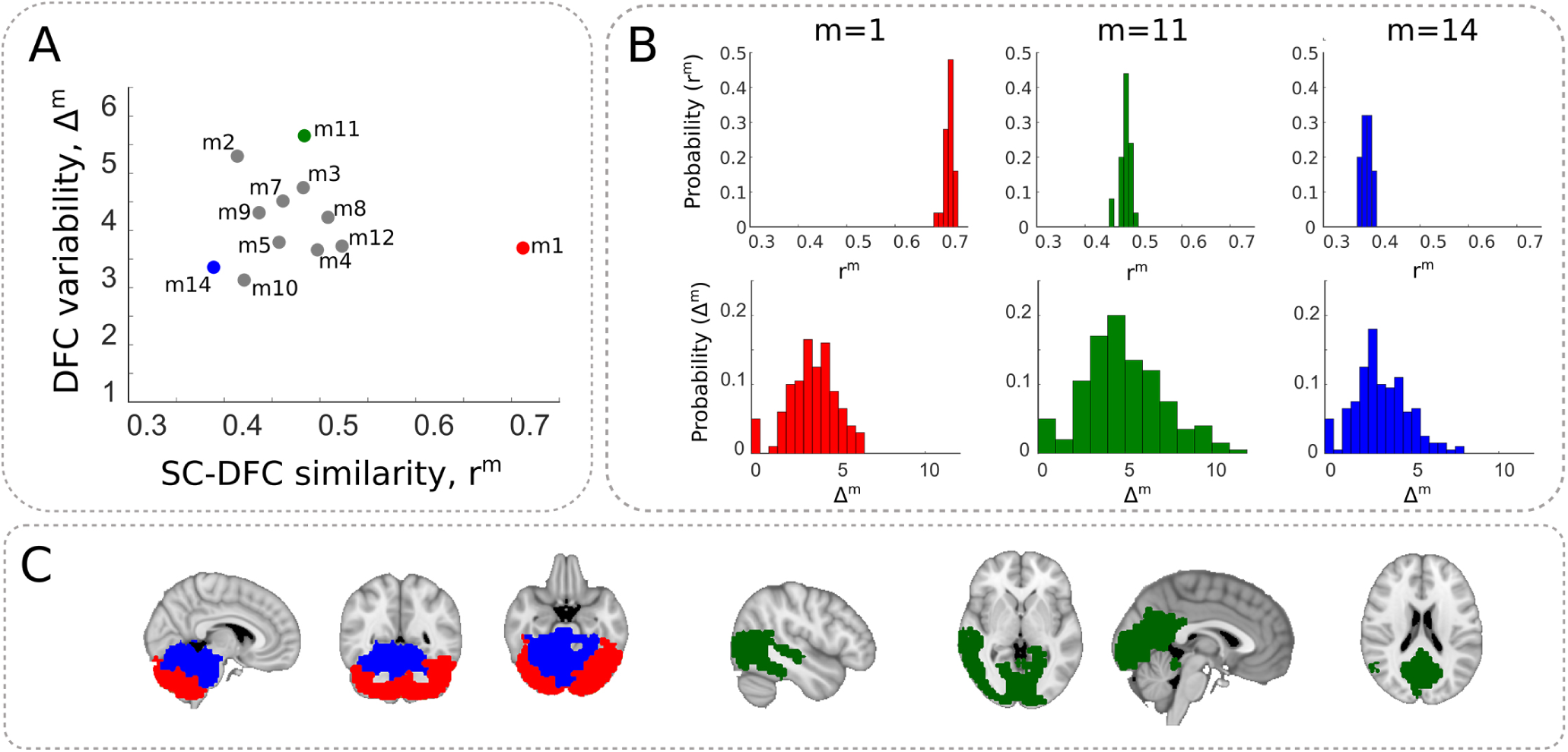
The amount of DFC variability and SC-DFC similarity define three outlier modules. **A**: For all SC structural modules, we plotted their representation in the plane (r^*m*^,Δ^*m*^), from which we detected three outlier cases: Module 1 (with the highest r^*m*^ value, red arrow), module 11 (with the highest Δ^*m*^ value, green arrow) and module 14 (with the lowest r^*m*^, blue arrow). The points represent the averaged values of r^*m*^ and Δ^*m*^ over all FC matrices corresponding to different time windows. **B**: Probability distribution of all values of r^*m*^ and Δ^*m*^, obtained at different windows. **C**: Anatomical representation of three modules. **A**,**B**,**C**: Results for a window length of *δ*=8, that for non-overlapping windows and a total number of 200 time points, it resulted in 25 different windows over which 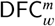 was calculated.

In addition to the SC-DFC similarity, we quantified the amount of DFC variability and found that module 11 –that is composed of several cortical structures, including the calcarine, middle and inferior temporal, lingual and precuneus (table 1)– exhibited the highest variability across time.

Figure 3A shows that, by looking to both metrics simultaneously, r^*m*^ and Δ^*m*^, both modules 1 and 14 had considerably smaller values of Δ^*m*^ than module 11, indicating that the cerebellar structures, as compared to others in the cerebrum, show much less DFC variability across time. Figure 3A represents mean values across windows and the histograms of possible values are shown in Figure 3B. The brain localization of these three modules is explicitly shown in Figure 3C.

Next, we asked whether our findings of the three modules obtained for a window length of *δ* = 8 were robust when varying the window over which DFC_*w*_ was calculated. In particular, we performed again all measurements for the following values of window lengths: *δ* = {4, 5, 7, 8, 10, 25}. Results are represented in Figure 4.

**Figure 4:**
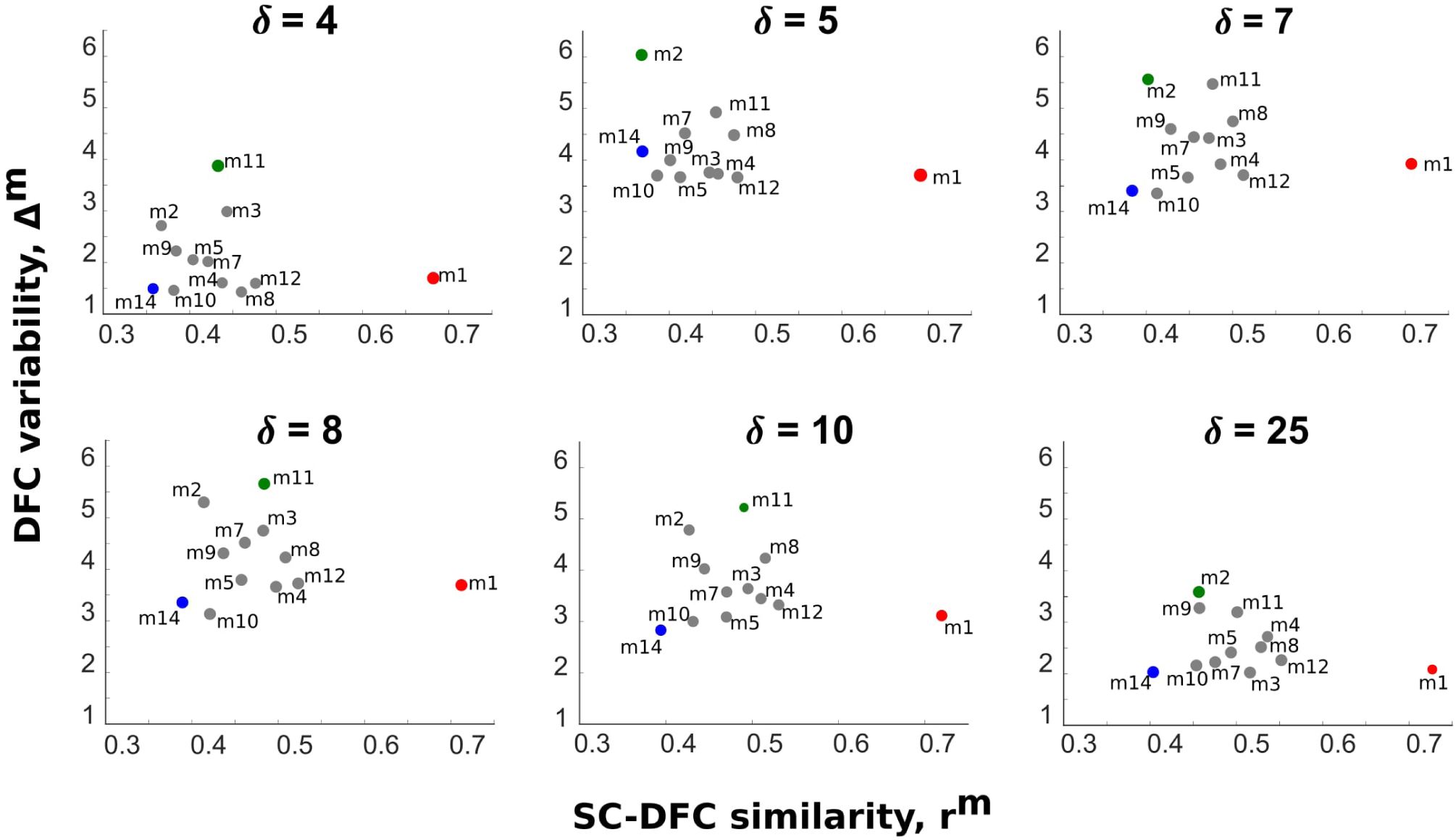
Robustness of the three relevant scenarios for different window lengths. The characteristics of modules 1 and 14, namely, to have respectively the highest and lowest value of r^*m*^ whilst keeping a low value of Δ^*m*^, preserved independently on the value of window length *δ*. However, the module 11, that in figure 3 had the highest value of Δ^*m*^, now this role switches between the module 2 and 11 when changing *δ*. Importantly, the two invariant modules 1 and 14 are both parts of the cerebellum.

Overall, the finding of three different outlier modules was preserved. In particular, modules 1 and 14, maintained their roles independently of the window length.

On the other hand, when looking along different window lengths at module 11 –that in figure 3 had the highest DFC variability– we noticed that this behavior was more variable across time windows, and modules 2 and 11 can interchange their position.

Finally, in addition to assess DFC by calculating the pairwise Pearson’s correlation between node time series, we also assessed DFC by taking into consideration synchronization metrics. In particular, by obtaining the Hilbert transform of the node time series *x*(*t*), and denoting it as 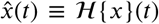, we calculated the complex analytical signal as 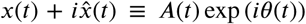, where *A*(*t*) and *θ*(*t*) are, respectively the instantaneous amplitude and the instantaneous phase of the analytical signal. Then, we obtained DFC_w_ matrices in two more different forms: (1) By calculating the pairwise Pearson’s correlation between time series of the instantaneous amplitude along different time windows, and (2) by calculating the pairwise Kuramoto order parameter, that for a window *w* of size *δ* is defined as 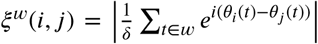, with | · | representing the modulus of a complex number. Our results using these metrics from the analytical signal (not shown here) are almost identical to those obtained in the main text relying on the Pearson’s correlation, measured directly from the time series; a similar equivalence between different ways to construct functional matrices has been also reported by other authors [38].

## Discussion

We have assessed the relation between SC and DFC at the level of structure-function modular organization using brain networks at a high spatial resolution, covering the entire brain with 2514 nodes of size equal of –on average– 20 voxels. Through modularity maximization of the SC population matrix, we obtained 14 non-overlapping structural modules that were used to reorder all functional matrices. This was done under the assumption that if segregated functions are associated with distinct structural modules, visualizing the DFC_*w*_ matrices in an ordering dictated by the structural one, might help to underline how SC constraints DFC_*w*_, a fundamental question that remains not fully understood.

Next, we analyzed the level of SC-DFC similarity in combination with the amount of DFC variability across time for all the previously identified modules, and characterized each module by the values of these two metrics. This allowed us to identify three extreme (or outlier) cases: 1. The presence of a fully cortical module that had the highest DFC variability; 2. The presence of a module at the posterior cerebellum which had the highest SC-DFC similarity while keeping low the DFC variability; 3. The presence of a module at the anterior cerebellum connected to the brain stem, which had the lowest SC-DFC similarity but also kept low the values of DFC variability.

Therefore, a common finding is that cerebellar networks have low variability in the DFC. The case of module 1, located at the posterior cerebellum, is fully consistent with another finding, namely, it has about twice more similarity between SC and DFC as compared to the rest of the modules, enabling the structure to constrain the dynamical connectivity. However, how module 14 with such a low value of SC-DFC similarity can keep the DFC variability low is more challenging to understand. On one side, in addition to the anterior part of the cerebellum, module 14 is composed of the brain stem which has vast connectivity to many other parts of the brain and body through major tracts such as the corticospinal, lemniscus, spinothalamic tracts [39]. Thus, by looking at the intra-module similarities between SC and DFC_*w*_, as we have done here, it is very possible that we ignored relevant connectivity aspects from this module to the rest of the brain, and for this reason, the measured value of intra-module SC-DFC similarity might be underestimated. Moreover, it is well known that the brain stem has a critical role in regulating sleep cycles, cardiac and respiratory functioning. Perhaps, such critical functions are not compatible with large DFC variability, as occurs in cortical networks, but to fully clarify this finding future research is needed.

It is important to emphasize that, in relation to the level of SC-DFC similarity, our unsupervised method has identified two distinct divisions within the cerebellum, the posterior and the anterior parts. This turns out to be a well-known and standard division of the cerebellum’s anatomy and functionality, shown in humans and animals [40]. Moreover, although the classical cerebellum division has grouped the lobules from 1 to 5 into the anterior part [41], our results include the lobule 6 in both the anterior and the posterior parts of the cerebellum, in agreement with task functional MRI studies, that found that cerebellar lobules 4-6 participated together in sensorimotor tasks [42].

When scrutinizing the brain areas to which modules 1 14 connect both structurally and functionally (figure 5), we found strong mutual connectivity between the anterior and posterior parts of the cerebellum (table 2), which is in agreement with previous work [40, 43]. Moreover, the anterior part projects also to the motor cortex among other areas as is also well established [41, 44, 45].

**Table 2.**
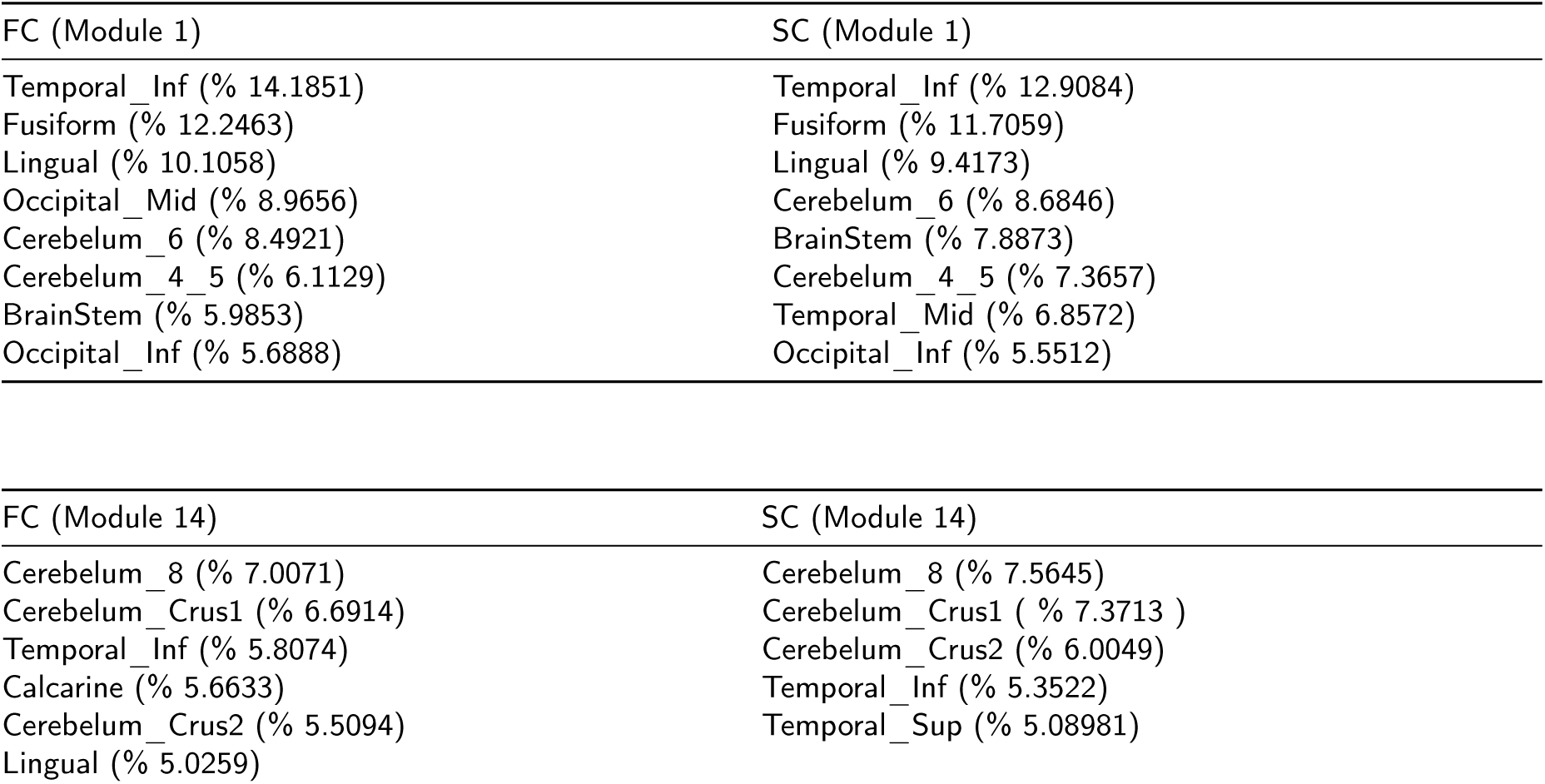
Functional and structural connectivity of modules 1 and 14 to rest of the brain. Notice that the two modules connect both functionally and structurally to one another. Module 14 also connected bilaterally to the anterior part of the paracentral lobule (2.65% of overlapping index), which is part of the supplementary motor cortex (see figure 5).

| FC (Module 1) | SC (Module 1) |
| --- | --- |
| Temporal_Inf (% 14.1851) | Temporal_Inf (% 12.9084) |
| Fusiform (% 12.2463) | Fusiform (% 11.7059) |
| Lingual (% 10.1058) | Lingual (% 9.4173) |
| Occipital_Mid (% 8.9656) | Cerebelum_6 (% 8.6846) |
| Cerebelum_6 (% 8.4921) | BrainStem (% 7.8873) |
| Cerebelum_4_5 (% 6.1129) | Cerebelum_4_5 (% 7.3657) |
| BrainStem (% 5.9853) | Temporal_Mid (% 6.8572) |
| Occipital_Inf (% 5.6888) | Occipital_Inf (% 5.5512) |

| FC (Module 14) | SC (Module 14) |
| --- | --- |
| Cerebelum_8 (% 7.0071) | Cerebelum_8 (% 7.5645) |
| Cerebelum_Crus1 (% 6.6914) | Cerebelum_Crus1 ( % 7.3713 ) |
| Temporal_Inf (% 5.8074) | Cerebelum_Crus2 (% 6.0049) |
| Calcarine (% 5.6633) | Temporal_Inf (% 5.3522) |
| Cerebelum_Crus2 (% 5.5094) | Temporal_Sup (% 5.08981) |
| Lingual (% 5.0259) |  |

<sup>1</sup>Data correction: slice-time correction, head-movement correction and intensity normalization
<sup>2</sup>Data filtering: regress out of confounding factors (movement time courses, average CSF signal and average WM signal) and bandpass filter

**Figure 5:**
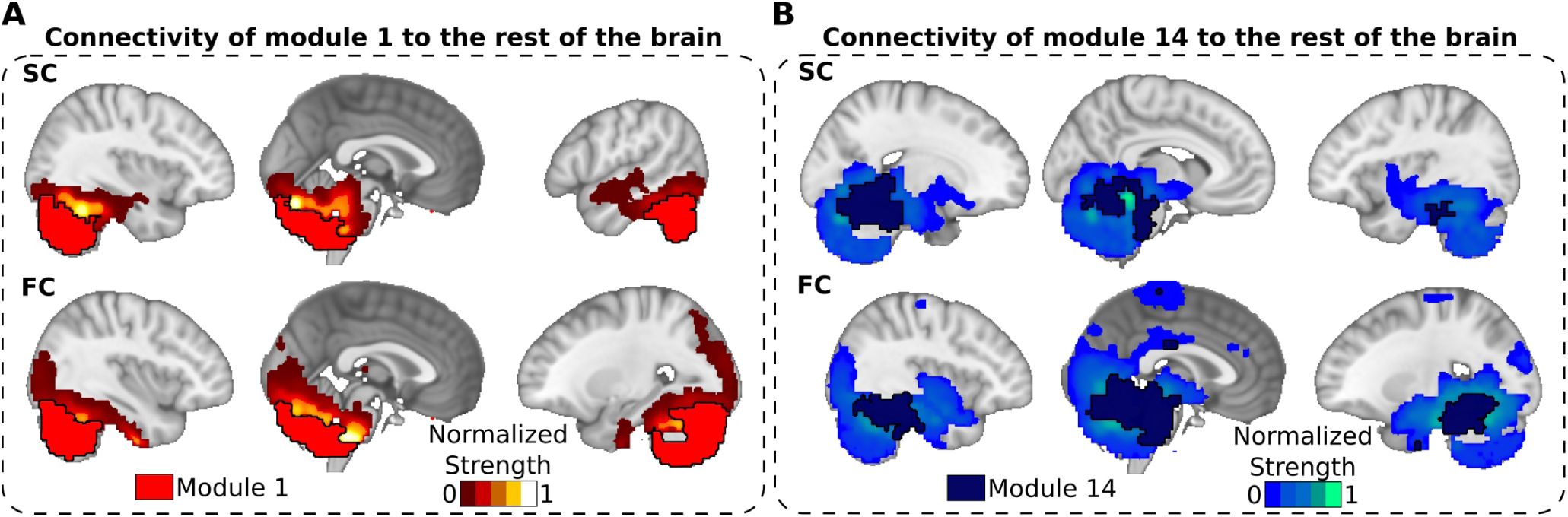
Structural and functional connections from the cerebellar networks to the rest of the brain. **A**: Structural (top row) and functional (bottom row) projections from module 1 (posterior part of the cerebellum, marked here with a black thick line surrounding the module) to the rest of the brain. **B**: Similar to panel A, but for module 14, formed by the anterior cerebellum and brain stem. **A**,**B**: Notice that both modules 1 and 14 strongly connect to one another (functionally and structurally), but module 14 also projects to motor cortex. Normalized strength accounts for the strength value of all connections arriving to a given node from all the nodes belonging respectively to the module 1 (red, panel A) and to the module 14 (blue, panel B). For FC, we only plotted values such that |FC| *>* 0.3.

To further understand the functional roles of modules 1 and 14, we projected both modules to a highly detailed cerebellar template [46, 47] and calculated their overlapping with the highly popular resting-state networks [48]. We found that module 1 overlapped with the default mode network (22.8%) and the frontoparietal network (19.2%), while module 14 did it with the somatomotor network (17.7%). Moreover, both modules overlapped similarly with the ventral attention network, with 13.3% of overlapping for module 1 and 10.5% for module 14. Thus, these results also show, in another way, that both cerebellar modules are differentiated but complementary, with the module 1 participating in high order cognitive networks and the module 14 in sensorial networks, but both of them sharing their participation in multimodal integration networks such as the ventral attention [49]. The overlapping between the module 1 and the frontoparietal network is also consistent with the fact that the cerebellum areas Crus 1 and 2 – included in module 1 – connect to the thalamus, and from here to prefrontal areas. This might explain why when a motor task is more difficult to perform, or simply it is a new task for the participant, posterior cerebellar activation shows up together with prefrontal activity to enhance cognitive monitoring of participant’s performance [42, 50–53].

Finally, what makes the cerebellar module 1 to have the dynamical functional connectivity constrained by the structural connectivity in such an extraordinary manner might indicate distinct operational and computational principles occurring in the cerebellum. Classically, cerebellar architecture has been modeled in a feedforward manner, in contrast with the highly recurrent circuits found in the cerebral cortices (see [54] and references therein), enabling the cerebellum to linearly integrate different inputs from other systems to generate outputs according to previously learned information patterns, following feedforward error-correction computations [55, 56]. Perhaps, such computational machinery makes the cerebellum’s information processing more reliable, in agreement with our results of low variability in its dynamical functional connectivity, but further research is needed to shed light on these findings.

In summary, our findings show that cerebellar networks have low variability of DFC as compared to other cerebrum networks. Cerebellar networks have extremely rich and complex anatomy and functionality [57], connecting to the brain-stem and cerebral hemispheres, and participating in a large variety of cognitive functions, such as movement coordination, bimanual coordination performance [58], executive function, visual-spatial cognition, language processing, and emotional regulation [59, 60]. But, as far as we know, the statement of low variability of the dynamical functional connectivity together with the strong similarity between their corresponding structural and functional networks of cerebellar networks has not been reported before.

## CRediT authorship contribution statement

**Izaro Fernandez-Iriondo:** Performed the analyses, Made the figures, Drafted the first manuscript, Wrote the manuscript. **Antonio Jimenez-Marin:** Performed the analysis, Made the figures, Wrote the manuscript. **Ibai Diez:** Preprocessed the images, Wrote the manuscript. **Paolo Bonifazi:** Wrote the manuscript. **Stephan P. Swinnen:** Acquired the data, Wrote the manuscript. **Miguel A. Muñoz:** Supervised the research, Equal last-author contribution, Wrote the manuscript. **Jesus M. Cortes:** Drafted the first manuscript, Supervised the research, Equal last-author contribution, Wrote the manuscript.

## Acknowledgements

A.J.M. acknowledges financial support from a predoctoral grant from the Basque Government (PRE_2019_1_0070). J.M.C. and P.B. acknowledge financial support from Ikerbasque (The Basque Foundation for Science) and from the Ministerio Economia, Industria y Competitividad (Spain) and FEDER (grant DPI2016-79874-R to J.M.C., grant SAF2015-69484-R to P.B.). J.M.C. acknowledges financial support from the Department of Economical Development and Infrastructure of the Basque Country (Elkartek Program, KK-2018/00032 and KK-2018/00090). S.P.S was supported by the the FWO Research Foundation Flanders (G089818N), the Excellence of Science funding competition (EOS; 30446199) and the KU Leuven Special Research Fund (grant C16/15/070). M.A.M acknowledges financial support from the Spanish Ministry and Agencia Estatal de investigación (AEI) through grant FIS2017-84256-P (European Regional Development Fund), as well as the Consejería de Conocimiento, Investigación y Universidad, Junta de Andalucía and European Regional Development Fund (ERDF), ref. SOMM17/6105/UGR for financial support.

---

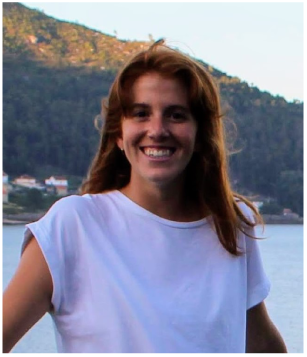

Izaro Fernandez-Iriondo received her Bachelor’s degree in Physics in 2019 from University of the Basque Country (UPV). She is currently studying a master’s degree in computer engineering and intelligent systems at the UPV in San Sebastian with the aim of finishing it with a machine learning project applied to structural and functional brain networks.

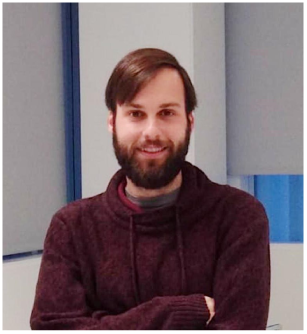

Antonio Jimenez-Marin is a PhD student of the Biomedical Research Program at the University of the Basque Country. His research is conducted in the Computational Neuroimaging Lab at Biocruces Bizkaia Health Research Institute. He obtained his degree in Telecommunication Technology Engineering at the University of Granada (2015) and his MsC in Biomedical Engineering at the University of the Basque Country (2018). His research interest is focused on brain connectivity in health and disease.

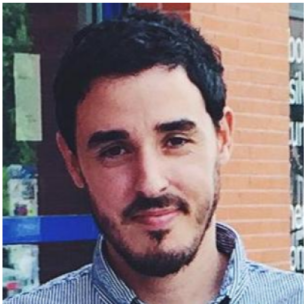

Ibai Diez received his Bachelor’s degree in Telecommunication Engineering in 2009 from Deusto University, Spain, and his PhD in 2015 from University of the Basque Country (UPV/EHU), Spain. He is currently a postdoctoral researcher in Neurology Department at Massachusetts General Hospital – Harvard Medical School. His research interest include structural and functional brain connectivity, biomedical data analysis and functional integration in the brain.

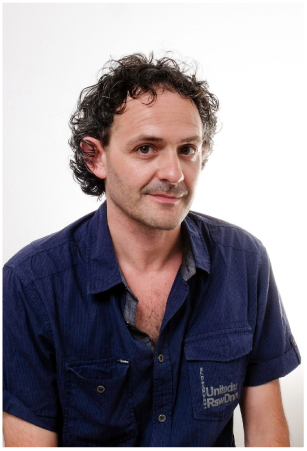

Paolo Bonifazi is a Tenured Ikerbasque Researcher at the Biocruces-Bizkaia Health Research Institute in Bilbao (Spain) within the Computational Neuroimaging group. He is a System Neuroscientist with a first degree in Physics (university of Perugia, Italy) and a PhD in Neuroscience (SISSA, Trieste, Italy). After two postdoctoral experiences in UK (with Prof. Hugh Robinson) and France (with Dr. Rosa Cossart), next he became Research Associate at the Tel Aviv University in the group of Prof. Ari Barzilai and late Prof. Eshel Ben-Jacob. His research interests are multi-disciplinarily focused on brain circuits’ structure and function, spanning from microcircuits to brain networks, bridging experimental and computational approaches, the latter inspired by complex networks.

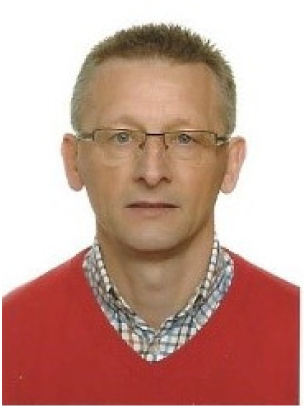

Stephan P. Swinnen started his research career in movement control at the University of California at Los Angeles (UCLA) (1983-85) under the direction of Prof. R. A. Schmidt. He completed his PhD at KU Leuven in 1987. He was awarded a Francqui Research Professor position (2013-2016) and currently directs the Movement Control & Neuroplasticity Research Group at KU Leuven (Belgium). His current research interest is focused on mechanisms underlying movement control and neuroplasticity in normal and pathological conditions using a multidisciplinary approach focusing on the study of brain function, structure, connectivity and neurochemicals.

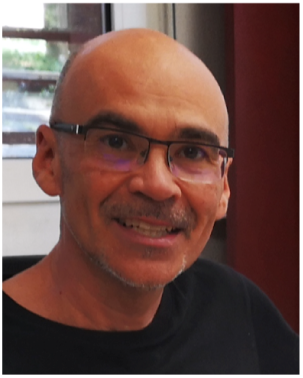

Miguel A. Muñoz is Full Professor in Physics at the University of Granada (Spain). He is an expert in statistical mechanics and has worked, among other issues, on non-equilibrium phase transitions, critical and collective phenomena and stochastic processes. In particular, he contributed to developing theory of “self-organized criticality” and helped developing approaches to the study of non-equilibrium phenomena, network theory and complex systems. His research interests spans from fundamental principles of statistical mechanics to interdisciplinary problems in evolutionary biology, theoretical ecology, and systems neuroscience.

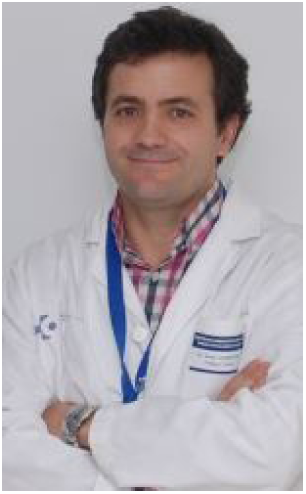

Jesus M. Cortes is an Ikerbasque Research Professor and the head of the Computational Neuroimaging Laboratory of the Biocruces-Bizkaia Health Research Institute in Bilbao (Spain). He teaches Brain Connectivity and Neuroimaging in the M.Sc of Biomedical Engineering. He obtained a Ph.D. in Physics in 2005 and performed three postdoctoral positions in The Netherlands (supervised by Prof. Bert Kappen), USA (supervised by Prof. Terry Sejnowski) and UK (supervised by Prof. Mark van Rossum). His area of research is now focused on brain connectivity, neuroimaging and machine learning methods applied to healthy and pathological conditions.

1 The alternative strategy, consisting in maximizing modularity in functional networks and using it to reorder the SC matrix was not used here.

